# From Alpha to Zeta: Identifying variants and subtypes of SARS-CoV-2 via clustering

**DOI:** 10.1101/2021.08.26.457874

**Authors:** Andrew Melnyk, Fatemeh Mohebbi, Sergey Knyazev, Bikram Sahoo, Roya Hosseini, Pavel Skums, Alex Zelikovsky, Murray Patterson

## Abstract

The availability of millions of SARS-CoV-2 sequences in public databases such as GISAID and EMBL-EBI (UK) allows a detailed study of the evolution, genomic diversity and dynamics of a virus like never before. Here we identify novel variants and sub-types of SARS-CoV-2 by clustering sequences in adapting methods originally designed for haplotyping intra-host viral populations. We asses our results using clustering entropy — the first time it has been used in this context.

Our clustering approach reaches lower entropies compared to other methods, and we are able to boost this even further through gap filling and Monte Carlo based entropy minimization. Moreover, our method clearly identifies the well-known Alpha variant in the UK and GISAID datasets, but is also able to detect the much less represented (< 1% of the sequences) Beta (South Africa), Epsilon (California), Gamma and Zeta (Brazil) variants in the GISAID dataset. Finally, we show that each variant identified has high selective fitness, based on the growth rate of its cluster over time. This demonstrates that our clustering approach is a viable alternative for detecting even rare subtypes in very large datasets.

## 1 Background

The novel coronavirus SARS-CoV-2, which is responsible for the Covid-19 disease, was first detected in Wuhan, China at the end of 2019 Wu *et al*. (2020); Zhou *et al*. (2020). Covid-19 was declared a global pandemic in March 2020 by the World Health Organization (WHO). According to recent data from the WHO (WHO), there have been almost 4 million deaths due to Covid-19, and there have been hundreds of millions of confirmed cases so far, while over 3 billion vaccine doses have been administered. As the virus continues to spread throughout countries and regions across the globe, it continues to mutate, as seen in the genomic variation among the millions of sequences which are available in public databases such as GISAID Elbe and Buckland-Merrett (2017). This mutational variability can be used to understand the evolution, genomic diversity and dynamics of SARS-CoV-2, and to generate hypotheses on how the virus has evolved and spread since it first originated.

An important part of these dynamics are the subsets of sequences (or subtypes) that vary more than others in terms of genomic content, which continue to emerge. In some cases, these subtypes appear in an atypically large number or are associated with an extremely high growth rate, indicating a possible fitness advantage (transmissibility, evasion from therapies or vaccines, etc.) of this genomic variation. The best example of this is the Alpha (or B.1.1.7 Rambaut *et al*. (2020)) variant, which differs from the typical sequence by about 30 mutations, and comprises hundreds of thousands of the currently available sequences. The Alpha variant was first detected in the UK at the end of summer 2020, where it grew to more than a third of the infected population in the UK by mid December 2020, as seen in the EMBL-EBI (UK) database EMBL-EBI (2020). One of the first variants of concern (VoCs), the Alpha variant has undergone much investigation, some studies Volz *et al*. (2021b) showing it to be between 40–80% more transmissible. This variant is now found in countries all over the world, some for which it is the dominant variant (*e*.*g*., The USA). Despite this, the origins of the Alpha variant are still contested, popular hypotheses including immunocompromised patients, the loss of records, or even minks as an intermediary host. There are now roughly a dozen variants identified around the globe (see Table 1) — an interesting question is whether they have or could potentially have the same degree of divergence as the Alpha variant.

**Table 1:**
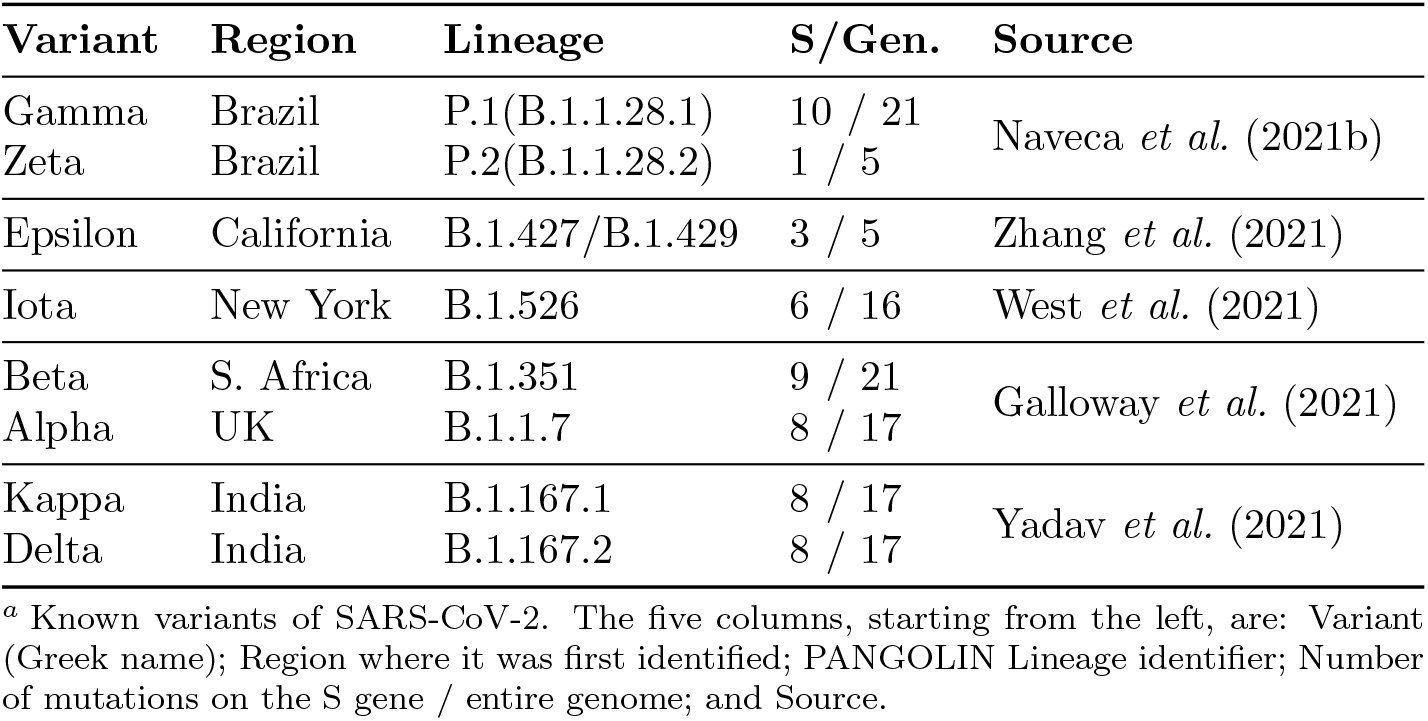
Some known variants of SARS-CoV-2^*a*^

The typical approach that is used to recover such knowledge from viral sequences is to construct a phylogenetic tree Hadfield *et al*. (2018); du Plessis *et al*. (2021) of evolution. However, with the high computational complexity of building a tree, more than a million sequences poses a scalability challenge for such methods Hadfield *et al*. (2018); du Plessis *et al*. (2021); Vrbik *et al*. (2015). An orthogonal approach to trees is to build transmission networks of infection — the structure of the network revealing general trends. In Skums *et al*. (2020), the authors show that such a network is *scale-free, i*.*e*., that few genomic variants are responsible for the majority of possible transmissions. A third alternative to studying the mutational variability of SARS-CoV-2 that we employ here is to *cluster* sets of sequences. While individual sequences are often unique, the sheer number of sequences available is expected to unveil meaningful groups and trends. Moreover, since most clustering techniques are much faster than, *e*.*g*., tree building Hadfield *et al*. (2018); du Plessis *et al*. (2021), such an approach can easily scale to the full size of the current datasets in order to leverage this information. The idea is that clusters of similar sequences should correspond to variants and subtypes, such as the Alpha variant mentioned above.

In this work, we cluster sequences by adapting methods which were originally designed for finding viral haplotypes from intra-host viral populations. The idea is that we use, *e*.*g*., CliqueSNV Knyazev *et al*. (2020), to find haplotypes in the massively inter-host viral population, using them as cluster centers in categorical clustering algorithms such as *k*-modes Huang (1997) in order to find subtypes. A measure we use to asses the clustering approaches, in the absence of a ground truth, is clustering entropy. This notion was introduced in Li *et al*. (2004), where the authors show that minimizing clustering entropy is equivalent to maximizing the likelihood that the set of sequences are generated from a set of subtypes, which closely models this setting of viral sequence evolution. Moreover, the authors of Li *et al*. (2004) show that clustering entropy is a convex function, allowing us to apply general optimization techniques such as the Monte Carlo method to minimizing entropy directly, as the objective. Finally, we use the subtypes found from our clustering techniques to patch gaps in the sequences, as an alternative to filling in the missing data with the reference genome, for example. This applies in particular to sequences collected before March, when SARS-CoV-2 sequencing and alignment were still being refined.

To validate our approaches, we use data from both GISAID and EMBL-EBI (UK) databases mentioned above (see Table 2). In a general comparison of methods and parameter settings on data from GISAID, we show that our CliqueSNV-based approach can achieve the lowest clustering entropy. What is interesting is that our gap filling approach allowed each method to lower its entropy even more. We then tested out a Monte Carlo based entropy minimization technique to show that it gives our method an even further edge on lowering the entropy. We compared various methods on their ability to find subtypes in the UK dataset, verifying this with the “ground truth” clusters which arise from metadata which tags each sequence with its lineage (*e*.*g*., B.1.1.7, the Alpha variant). Our CliqueSNV-based approach identified the Alpha variant with significantly higher precision and specificity compared to other methods, based on these metadata. This reinforces the notion that clustering entropy is an appropriate measure of the quality of clustering in this context. We then used our CliqueSNV-based clustering approach to identify subtypes in the GISAID dataset, again verifying this from metadata. While our method clearly identified the Alpha variant, it was also able to detect the lesser represented Beta (South Africa), Epsilon (California), Gamma and Zeta (Brazil) variants with specificities around 50%. What is particularly interesting about this is that these lesser represented variants comprise less than 1% of the GISAID dataset (which contains more than one million sequences), yet our method was able to detect them with a much higher specificity. Finally, we validate our approaches to finding subtypes using the *fitness coefficient*, a third measure of clustering quality which is orthogonal to both entropy and specificity mentioned above. The fitness coefficient, introduced in Skums *et al*. (2012), is an assessment of the selective fitness of a subtype, based on the number of sequences in the corresponding cluster, and the rate at which this grows over time. Our results show that fitness tends to corroborate with these other two measures, further strengthening our results. This demonstrates that we can use clustering for the identification and surveillance of new variants, which have the potential to grow quickly, or become a threat to public health.

**Table 2:** Datasets used in the experiments^*a*^

| Dataset | Database | Start | End | No. Sequences |
| --- | --- | --- | --- | --- |
| GISAID 1 | GISAID | 2019-12-24 | 2020-11-07 | 209 334 |
| GISAID 1A | GISAID | 2019-12-24 | 2020-03-05 | 3 688 |
| UK | EMBL-EBI | 2020-01-29 | 2020-12-29 | 88 008 |
| GISAID 2 | GISAID | 2019-12-24 | 2021-04-04 | 1 000 982 |

<sup>a</sup> The four datasets that are used in the experiments of Sec. 5. The five columns, starting from the left, are: Name we use here; Database it is from (GISAID Elbe and Buckland-Merrett (2017) or EMBL-EBI EMBL-EBI (2020)); Earliest collection date of any sequence; Latest collection date; and Number of sequences. Note that GISAID 1A is a subset of GISAID 1, which is, in turn, a subset of GISAID 2.

This paper is structured as follows. In Sec. 2, we detail the various approaches to clustering, gap filling and entropy minimization that we use in this work. In Sec. 3, we specify the measures that we used to assess the clustering approaches — clustering entropy and the fitness coefficient. In Sec. 4, we give details on the datasets that we use in this study, as well as some of the known variants which we can expect to find in their metadata. In Sec. 5, we report the results of the experiments we performed to assess the various clustering approaches, gap filling, entropy minimization, and the identification of subtypes. Sec. 6 then concludes the paper with a discussion of the contributions of our approach, in light of these results.

## 2 Methods used in clustering

We outline in this section all of the methods that were used in clustering nucleotide sequences of the SARS-CoV-2 virus.

### 2.1 CliqueSNV

Since we are clustering viral sequences in order to identify subtypes, we propose to use currently existing tools that were developed to identify subtypes in intra-host viral populations from NGS data reviewed in Knyazev *et al*. (2021), such as Savage Baaijens *et al*. (2017), PredictHaplo Prabhakaran *et al*. (2014), aBayesQR Ahn and Vikalo (2018), *etc*. However, our setting is slightly different, where the data consists of large collections of *inter-host* consensus sequences gathered from different regions and countries around the world Elbe and Buckland-Merrett (2017); EMBL-EBI (2020). We expect, however, that such tools are appropriate at this scale: now the “host” is an entire region or country, and we reconstruct the subtypes, or variants, and their dynamics within these regions or countries. The SARS-CoV-2 sequences in GISAID are consensus sequences of approximate length 30K. Such sequences by quality and length have similar properties as PacBio reads. We choose CliqueSNV since it performed very well on PacBio reads Knyazev *et al*. (2020).

### 2.2 *k*-modes

Since nucleotide sequences can be viewed as objects on *categorical* attributes — the attributes are the genomic sites, and the categories are A, C, G, T (and –, a gap) — we use *k*-modes Huang (1997, 1998) for clustering. The *k*-modes approach is almost identical to *k*-means Anderberg (1973); MacQueen *et al*. (1967), but it is based on the notion of *mode* (rather than Euclidean mean), making it appropriate for clustering categorical data. Indeed, the Euclidean mean of three nucleotides has little meaning in this context, and may not even be well-defined, *e.g*., in cases where the “distance” from A to G is different than from G to A. Similar observations were made in the context cancer mutation profiles Ciccolella *et al*. (2020), in the form of absence/presence information. Treating these as categories, in using *k*-modes (rather than as 0’s and 1’s, in using *k*-means) resulted in a clustering approach Ciccolella *et al*. (2021a) that, when used as a preprocessing step, allowed cancer phylogeny building methods to attain a higher accuracy Ciccolella *et al*. (2021b), and in some cases with much lower runtimes Jahn *et al*. (2016). We briefly describe the *k*-modes approach in the context of clustering nucleotide sequences as follows.

The *mode q* of a cluster *C* of sequences on categorical attributes *𝒜* = {A, C, G, T, –} is another “sequence” on *𝒜* which minimizes

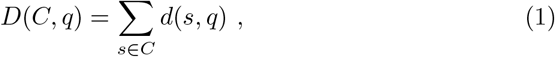

where *d* is some categorical dissimilarity measure (*e.g*., Hamming distance) between the sequences we are considering. Note that *q* is not necessarily an element of *C*. For a set *S* of sequences on attributes *𝒜*, we are given some initial set *Q* = {*q*_1_, …, *q*_*k*_} of *k* cluster “centers” (each on 𝒜). The *k*-modes approach (similarly to *k*-means) then operates according to the iteration:

- compute the dissimilarity *d*(*s, q*) between each sequence *s* ∈ *S* and each center *q* ∈ *Q*;
- assign each sequence *s* ∈ *S* to the closest center based on the first step, resulting in a clustering with *k* clusters; and
- compute the mode of each cluster from the second step, resulting in a new set *Q*′ of *k* centers;

until convergence, *i.e*., the clustering does not change after an iteration.

In this paper we cluster sequences of SARS-CoV-2 with *k*-modes using three different ways to compute the initial set *Q* of cluster centers, and using two different dissimilarity measures *d*. The three ways to compute the initial set of cluster centers are:

1. choose *k* random sequences from the dataset;
2. choose *k* centers that are maximally pairwise distant from each other; and
3. use the centers (the subtypes) found by CliqueSNV.

The two different dissimilarity measures that we use are (a) the Hamming distance, and (b) the TN-93 distance Tamura and Nei (1993).

### 2.3 MeShClust

For comparison purposes, we also apply methods designed for clustering metagenomics and multiviral sequencing data. We clustered the sequences using MeShClust James *et al*. (2018), an unsupervised machine-learning method that aims to provide highly-accurate clustering without the need for user-specified similarity parameters (these are learned).

However, this approach is intended for use with data sets containing genomes of multiple different viruses. In particular, it was validated on a data set containing 96 sequences of average length of 3K–12K, coming from 9 different viruses. On the other hand, SARS-CoV-2 data sets usually contain several hundred thousand sequences of a single virus, with genome length averaging around 30K.

### 2.4 Monte-Carlo Based Entropy Minimization

We use clustering entropy Li *et al*. (2004) to assess the various clustering methods that we propose in this work (see Sec. 3.1). For this reason, we also employ a technique aimed directly at minimizing clustering entropy as the objective. We first define clustering entropy in the following.

Formally, we have a set *S* of aligned nucleotide sequences on the set *X* of genomic sites. Since they are aligned, sequences can be viewed as rows of a matrix and, when restricted to a site *x* ∈ *X*, can be viewed as columns of this matrix. Let *𝒩* = {A, C, G, T} be the four nucleotides, not counting the gap (–) character. Using the notation of Li *et al*. (2004), the entropy *Ĥ*_*x*_(*C*) of a subset *C* (a cluster) of sequences from *S* at site *x* ∈ *X* is then

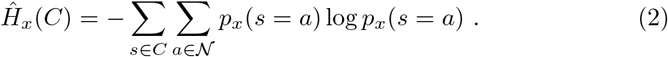

Note that *p*_*x*_(*s* = *a*) — the probability that a sequence *s* ∈ *C* has nucleotide *a* at site *x* — essentially amounts to the relative *frequency* of nucleotide *a* ∈ *𝒩* in *C* at site *x*. The entropy *Ĥ*_*X*_ (*C*) of subset *C* of sequences on a subset *X* of sites is then

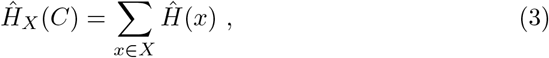

that is, we simply sum up the entropies at the individual sites. Since the set of sites will always correspond to the SNV sites of our sequences, we will use simply *Ĥ* (*C*) for the entropy of a subset (a cluster) of sequences from hereon in. The *expected* entropy Li *et al*. (2004) of a clustering ℂ = *C*_1_, …, *C*_*k*_ of sequences is then

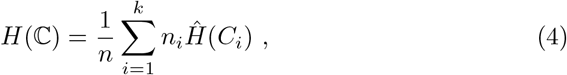

where *n*_*i*_ = |*C*_*i*_|, the number of elements in cluster *C*_*i*_, and *n* is the total number of sequences. For completeness, the *total* entropy of a clustering is simply the sum

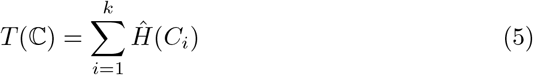

of the individual entropies of each cluster (not weighted by *n*_*i*_).

In Li *et al*. (2004), the authors prove that the entropy Eq. 4 is a convex function, allowing any optimization procedure to reach a global minimum. It is because of this property that we can use techniques aimed directly at minimizing clustering entropy as the objective. The Monte-Carlo method is broad class of computational algorithms that rely on repeated random sampling to optimize some criterion. In this context, we are randomly sampling clusterings of sequences in order to minimize Eq. 4. The basic idea is that we start with some clustering — note that the clustering corresponding to placing all sequences in the same cluster has maximum entropy, by definition. The Monte-Carlo process then operates according to the iteration:

- from the current clustering, randomly pick a sequence from some cluster and place it into another cluster, resulting in a new clustering;
- compute the entropy (Eq. 4) of the new clustering; and
- accept this new clustering, if the entropy has decreased, otherwise keep the current clustering;

until convergence, *i.e*., the clustering does not change after some number *θ* of iterations.

In Li *et al*. (2004), the authors prove the concept of applying the Monte-Carlo method to entropy minimization by implementing a very basic procedure similar to the above, and then demonstrate it on a small dataset. Since our datasets are on a much larger scale (millions of sequences on 30K genomic sites), the basic iteration which randomly samples a single sequence in each iteration would need many iterations for a very small improvement. For this reason we apply the following preprocessing step, to improve the convergence. Rather than using all (30K) columns, we first sort the columns according to their (unclustered) entropy value. We then select the *n* columns, or *tags*, with highest entropy. Next, we then run the above Monte-Carlo process on the reduced dataset with the *n* tags. This results in a clustering (of the rows), to which we then apply to the original set of all columns.

### 2.5 Filling gaps

Finally, the set of SARS-CoV-2 sequences that we deal with contain missing nucleotides, due to gaps or deletions. This is particularly true with GISAID sequences collected from December 2019 to the end of March 2020, when sequencing, alignment, *etc*., were less refined. This is further complicated by the presence of deletions, which could be confused with gaps.

Here, we attempt to use the clustering obtained by some clustering method in order to fill the gaps. That is, rather than uniformly filling all sequences with, *e.g*., the reference genome, we fill each sequence with the center of its cluster. The idea is that if a clustering performs well, then the sequences of a cluster should correspond to a subtype. In this case, the center — a consensus sequence of this subtype — should be much closer to any sequence of its cluster than the reference genome, resulting in a more accurate filling of the gaps.

## 3 Measures for assessing clustering quality

In this section, we present two measures for assessing clustering quality, in order to compare the various clustering methods that we outlined in the previous section. The first measure is clustering entropy, an internal evaluation criterion that reflects the underlying processes which generate a set of viral sequences. The second is a measure of the selective fitness of clusters, based on how their rates of change in size vary over time.

### 3.1 Clustering entropy

Since we are comparing various clustering methods without knowing a ground truth, we need to consider an *internal* evaluation criteria. Many of the commonly used criteria require some notion of distance, or dissimilarity measure, between the objects being clustered. For example, criteria such as the Calinski-Harabasz Index Caliński and Harabasz (1974) or the Gap Statistic Tibshirani *et al*. (2001) rely on the Euclidean distance, while the Davies-Bouldin Index Davies and Bouldin (1979) or the Silhouette Coefficient Rousseeuw (1987) require this distance (or dissimilarity) to be a *metric*. For the same reason that we use *k*-modes for clustering — sequences are objects on categorical attributes which take values A, C, G, T (and –, a gap) — criteria based on the Euclidean distance are not appropriate. Moreover, because the various dissimilarity measures that we use within the *k*-modes framework for clustering are not Euclidean (Hamming distance), or even a metric (TN-93 distance, see Table 1 of Tamura and Nei (1993)), even criteria such as the Davies-Bouldin Index or Silhouette Coefficient would not apply.

The clustering entropy Li *et al*. (2004) (Eq. 4 and Eq. 5) is an internal evaluation criterion that was shown to generalize any distance-based criterion, and does not even require any notion of distance or dissimilarity. Hence, for the reasons mentioned above, the clustering entropy criterion is appropriate in our case. Moreover, clustering entropy naturally reflects the fact that the population of viral sequences comes from a number of subtypes. Clustering entropy can be formally derived using a likelihood principle based on Bernoulli mixture models. In these mixture models, the observed data are thought of as coming from a number of different latent classes. In Li *et al*. (2004), the authors prove that minimizing clustering entropy is equivalent to maximizing the likelihood that set of objects are generated from a set of (*k*) classes. This reflects the underlying processes which generate a set of viral sequences: that they evolved from a set of (*k*) subtypes.

This relates closely to the widely-used notion of *sequence logo* Schneider and Stephens (1990): a graphical representation of a set of aligned sequences which conveys at each position both the relative frequency of each base (or residue), and the amount of information (the entropy) in bits. A clustering of viral sequences of low entropy then relates to a reliable set of sequence logos (in terms of information), and can hence shed light on the possible biological function of the viral subtype that each such logo (or related motif) represents.

### 3.2 Fitness

We use a mathematical model proposed in Skums *et al*. (2012) for the calculation of a numerical measure of the *fitness* of a quasispecies. This model is used here to calculate the fitness of a cluster, based on how the rate of change in size (number of sequences it contains) varies over time. For a set *C*_1_, …, *C*_*k*_ of clusters, *X*_*i*_(*t*) denotes the size of cluster *C*_*i*_ at a particular time *t*. The fitness coefficient is calculated using *h*_*i*_, which is the cumulative sum of the *X*_*i*_. It follows that 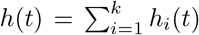 is the total infected population size at time *t*. Each *h*_*i*_(*t*) is normalized over *h*(*t*), which is denoted by *u*_*i*_(*t*), that is,

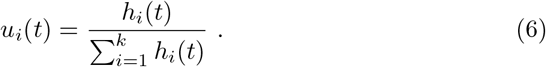

Using cubic splines, *u*_*i*_(*t*) and *h*(*t*) are interpolated over the time period and the derivatives 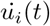 and 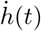 are calculated. The *fitness function g*_*i*_, for each cluster *C*_*i*_ is then defined as

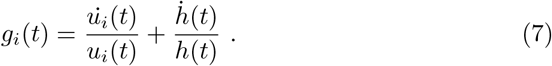

The *fitness coefficient r*_*i*_, which is the average fitness over the time period *T* (composed of the times *t*) for cluster *C*_*i*_ is then

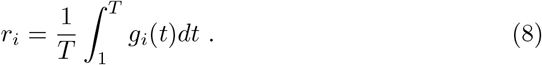

In order to reduce sampling error, we use the Poisson distribution to draw random samples. For each cluster at time *t*, a sufficiently large number of random samples is drawn from the Poisson distribution on *X*_*i*_(*t*) as the expectation of the interval. Then *X*_*i*_(*t*) is replaced by the mean value of these random samples. This is repeated a sufficiently large number of times (*e.g*., 100) to calculate a set of Poisson-distributed sizes. The fitness coefficient calculation is then applied on each repetition separately and a confidence interval of this fitness coefficient is obtained.

## 4 Datasets

In this section we outline the datasets that we used in the experiments of the next section. We first give a brief overview of well-known subtypes, or *variants*, from the literature, and then describe the three datasets we use, which are known to contain different proportions of these variants.

### 4.1 Known variants

Since its emergence in November 2019 Deasy *et al*. (2020), SARS-CoV-2 has evolved into different variants. Divergences in mutation at the genomic level have been observed in different regions of the world as new infectious variants are emerging. The following is a description of some of the well-known variants to date. A more complete list can be found in Table 1.

#### 4.1.1 Alpha variant (UK)

The Alpha variant, also known as the B.1.1.7 variant of SARS-CoV-2 was first identified in Kent, UK, in late summer to early autumn 2020. It has the highest transmissibility of any lineage, with a 50% to 100% reproduction rate Volz *et al*. (2021a). The first case was reported on December 14, 2020, and this variant is now detected in over 30 countries, with more than 15 thousand people affected worldwide Galloway *et al*. (2021). Of the many genomic mutations that characterize this variant, it has a 69/70 deletion and a mutation at position 501, which affects the conformation of the receptor binding domain (RBD) of the spike protein of SARS-CoV-2. It has 17 mutations which include 14 amino acids and 3 in-frame deletions at open-reading frame (ORF) 1 a/b, ORF 8, spike (S), and N gene regions. These mutations have biological implications and have resulted in diagnostic failures Ramírez *et al*. (2021).

#### 4.1.2 Beta variant (South Africa)

The first case of the Beta variant, also known as B.1.351, was identified in Nelson Mandela Bay, South Africa, in October 2020. This lineage was predominant by the end of November 2020 in the Eastern and Western Cape Provinces of South Africa. By January 2021, there were 415 known cases of infection with this variant, found in 13 different countries. This variant has eight mutations in the S gene region, including three mutations SK417N, E484K and N501Y that affect the RBD of the spike protein. These three mutations can be the reason for increased transmissibility, and can also lead to alterations in conformation that could pose a challenge for the effectiveness of vaccines Galloway *et al*. (2021); Zucman *et al*. (2021); Tang *et al*. (2021).

#### 4.1.3 Gamma and Zeta variants (Brazil)

The Gamma variant, also known as P.1(B.1.1.28.1), was initially identified in February 2020, in Japanese travelers coming from Amazonas State, Brazil. It was first reported in a 29-year-old female resident of Amazonas State. The P.1 lineage has mutations K417T, E484K, and N501Y in the S gene region, which affect the RBD of the spike protein. The Zeta variant, also known as P.2(B.1.1.28.2) was first identified in Rio de Janeiro, Brazil. It shares the mutation E484K with the Gamma variant Naveca *et al*. (2021a).

#### 4.1.4 Epsilon variants (California, USA)

In July 2020, the first case of the Epsilon variants, also known as the CAL.20C or B.1.427/B.1.429 variants of SARS-CoV-2, was identified in Los Angeles County, California, USA. The Cedars-Sinai Medical Center (CSMC) reported that the second B.1.429 Epsilon variant contains five mutations at ORF 1 a (I4205V), ORF 1 b (D1183Y), and S gene mutations S13I, W152C and L452R. Mutation L452R is correlated with higher infectivity Zhang *et al*. (2021). The Epsilon variants are spreading in the US and in 29 other countries McCallum *et al*. (2021).

#### 4.1.5 Iota Variant (New York, USA)

The Iota variant, also known as B.1.526, was first found in November 2020 in New York, USA. At that time, the number of sequences of the Iota variant comprised less than 1% of all sequences in the GISAID database Elbe and Buckland-Merrett (2017). Scientists from Caltech noticed a surge in growth of this number by roughly one third by February 2021. This variant has mutations L5F, T95I, D253G, E484K or S477N, D614G, and A701V in the S gene region — mutations E484K and S477N affecting the RBD of the spike protein. Note that the E484K mutation causes attenuation in *in vitro* neutralization, and is found in other variants of concern (VOCs) Thompson *et al*. (2021); West *et al*. (2021), such as the Beta, Gamma and Zeta variants, described above.

### 4.2 Datasets used

In our experiments, we use four different datasets, three of which are various snapshots of the GISAID Elbe and Buckland-Merrett (2017) database at different time points, and the fourth is a dataset obtained from the EMBL-EBI EMBL-EBI (2020) database in the UK. These datasets are summarized in Table 2, and then each one is explained in more detail in its corresponding subsection below.

#### 4.2.1 GISAID 1

The first dataset consists of sequences submitted to the GISAID Elbe and Buckland-Merrett (2017) database up until November 2020. This dataset contains sequences from all over the world. Since this dataset covers the period of time from December 2019 to March 2020, some of these sequences have a sizeable number of gaps.

#### 4.2.2 GISAID 1A

The second dataset consists of sequences submitted to GISAID up until the beginning of March 2020. This smaller dataset, a subset of GISAID 1, was designed in order to test out the Monte Carlo optimization procedure described in Sec. 2.4.

#### 4.2.3 UK

The third data set consists of sequences submitted to the EMBL-EBI EMBL-EBI (2020) database from the end of January 2020 to the end of December 2020. Since this database is in the UK, and given the collection period, this dataset contains a sizeable portion of the Alpha variant.

#### 4.2.4 GISAID 2

The third data set consists of all sequences submitted to GISAID up until April 2021. Since many of the known variants mentioned above have been well-documented by April 2021, this dataset contains a sizable portion of sequences annotated as being from the Alpha, Beta, Gamma, Epsilon and Zeta variants. Such labels correspond to “ground truth clusters” for which we can compute the precision, specificity, *F*_1_ score, etc., of a clustering obtained with a given method.

## 5 Experimental results

In this section we report the results of our approaches of clustering and gap filling using the four data sets mentioned in Sec. 4.2, above. For all datasets, we align the sequences and trim the first and last 50bp of the aligned sequences. We use default parameters for running CliqueSNV to find initial cluster centers, in all cases setting the minimum cluster frequency to be at least 1% of the population. We refer to the approach of using CliqueSNV to find the initial centers, followed by clustering with *k*-modes as our CliqueSNV-based approach (setting 3. of Sec. 2.2).

These experiments, and their results, are grouped as follows: a general comparison of clustering and gap filling approaches, using the GISAID 1 dataset (Sec. 5.1); a test of our Monte-Carlo based entropy minimization procedure introduced in Sec. 2.4, using the GISAID 1A dataset (Sec. 5.2); and a demonstration of the use of various clustering methods for finding subtypes, using the UK and GISAID 2 datasets (Sec. 5.3).

### 5.1 Comparison of clustering approaches

Using the GISAID 1 dataset, our CliqueSNV-based approach identified at most 66 subtypes (smallest *k* which achieves minimum cluster frequency *≥* 1%), which vary in proportion between December 2019 and November 2020. We report the relative distributions over time of these different subtypes in Fig. 1 and Fig. 2, in a similar way to that of Fig. 3 of du Plessis *et al*. (2021).

**Figure 1:**
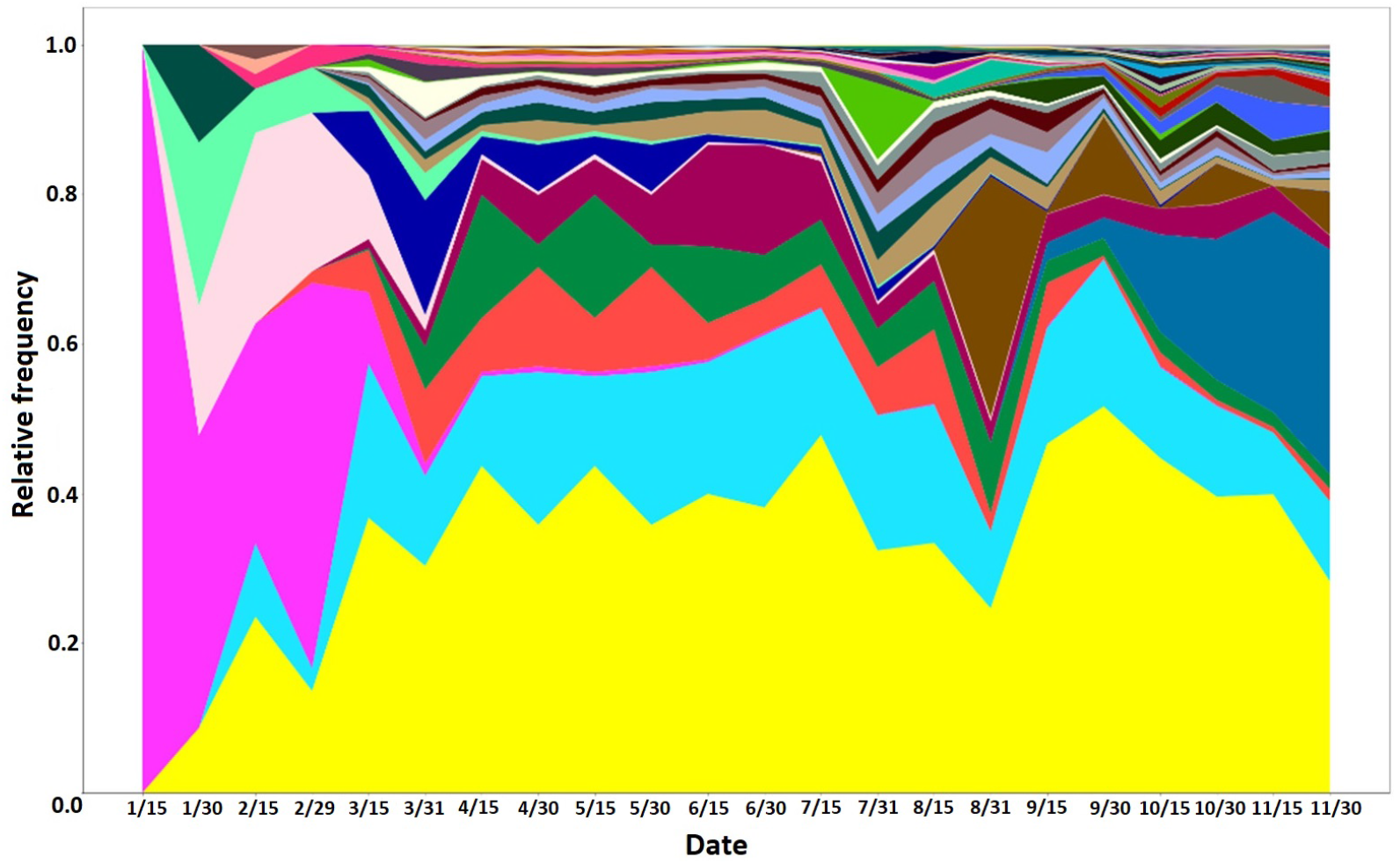
Subtype distribution (GISAID dataset, 15-day window, relative count)

**Figure 2:**
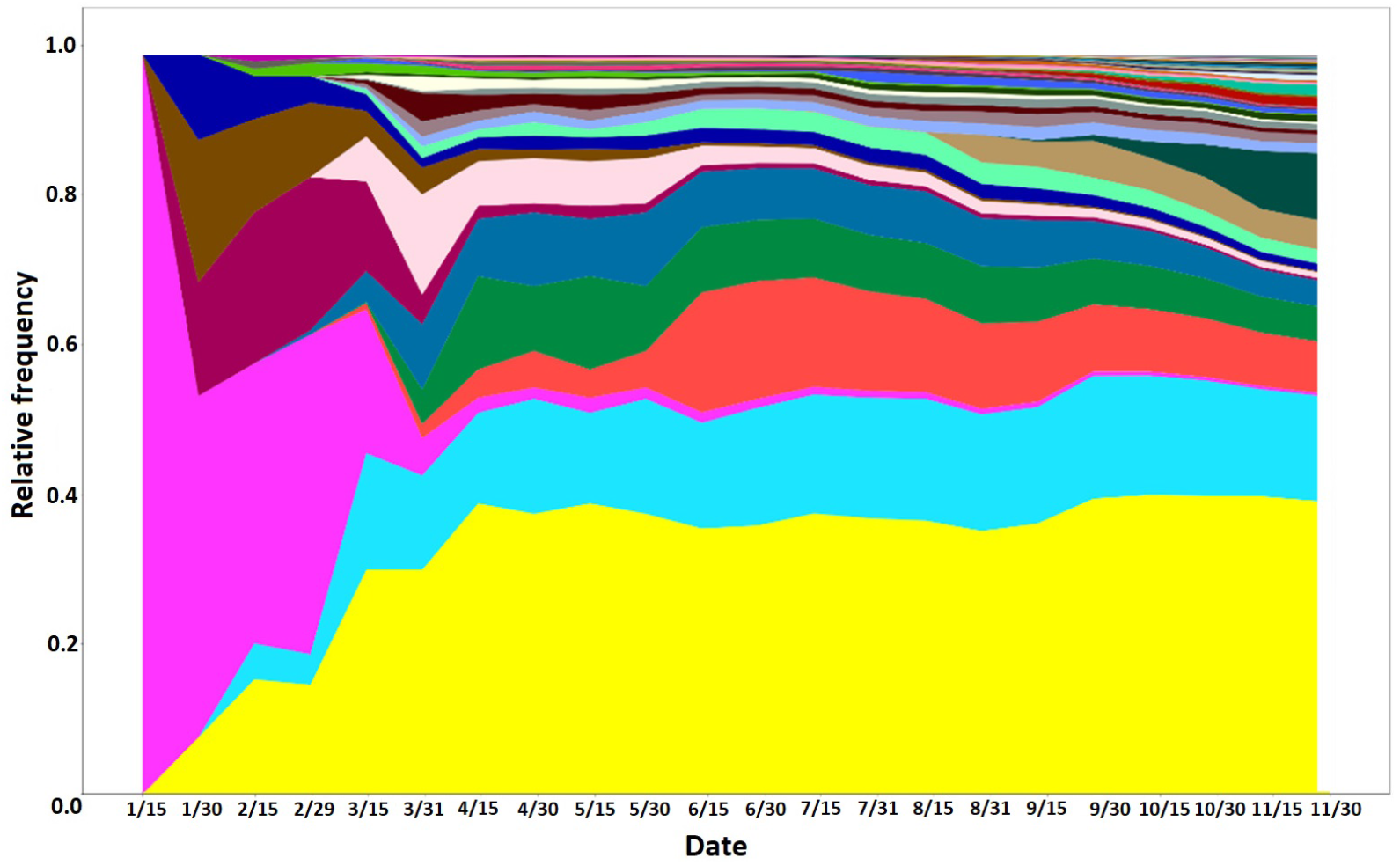
Subtype distribution (GISAID dataset, cumulative, relative count).

**Figure 3:**
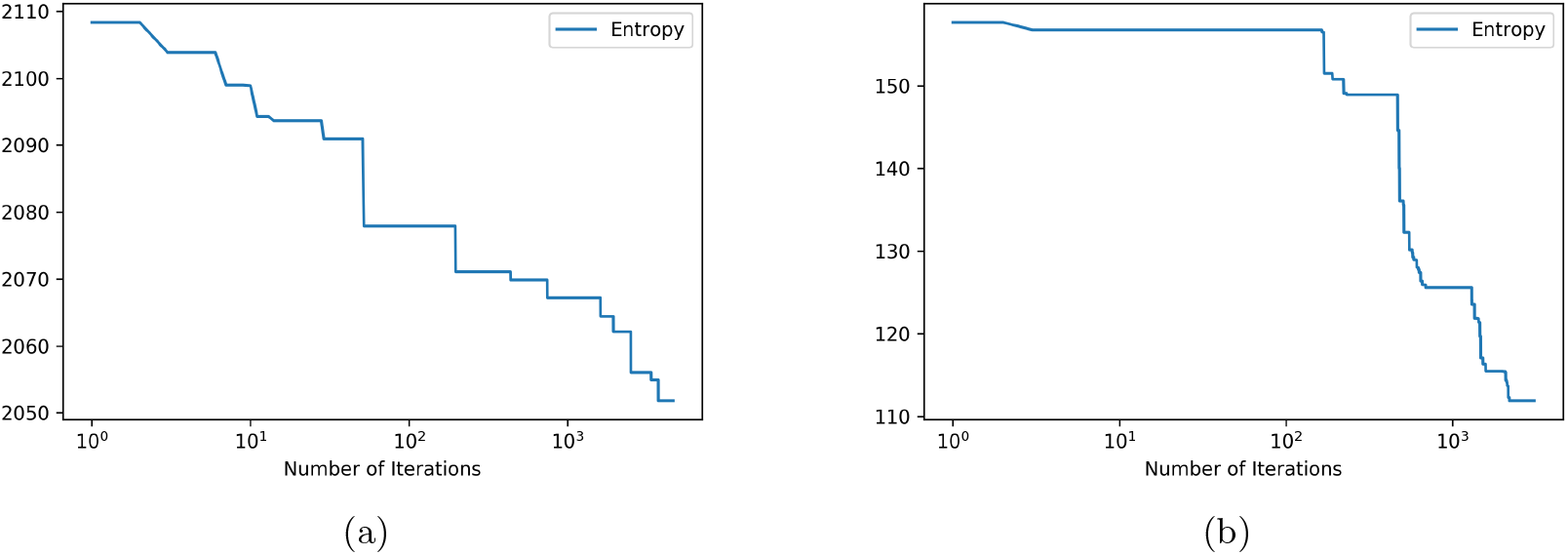
The entropy descent of our Monte-Carlo method applied to the initial clustering obtained by CliqueSNV-based clustering of the GISAID 1A dataset after having preprocessed to: (a) *n* = 28000 tags; and (b) *n* = 1000 tags. Note that in the latter table, the entropy is in terms of just the 1000 tags — the optimal clustering in terms of these 1000 tags then applied to the original set of all columns, for the final entropy 945.9 seen in Table 6. Note that a threshold of *θ* = 1000 (see Sec. 2.4) was used in both cases.

Table 3 gives an assessment of all clusterings (and gap fillings) of the GISAID 1 dataset that were computed for the various settings mentioned in Sec. 2.2, in terms of both the expected entropy (Eq. 4) and total entropy (Eq. 5). While any form of clustering achieves a better expected (and total) entropy than not clustering at all, our CliqueSNV-based approach tends to outperform all other forms of clustering using either Hamming or TN-93 distance. This demonstrates that CliqueSNV finds meaningful centers in these inter-host viral data. Based on these results, from hereon in we use only the Hamming distance setting of our CliqueSNV-based clustering (setting 3.(a) of Sec. 2.2, second-last line of Table 3), and the random centers initialization and Hamming distance setting (setting 1.(a) of Sec. 2.2, second line of Table 3) of *k*-modes, unless otherwise indicated. Finally, by filling gaps in sequences based on the center of its cluster, we achieve an even lower expected (and total) entropy. This highlights the value of a cluster-based approach for filling gaps. For example, the entropy of the dataset without clustering remained high even after filling gaps, which would, by definition, be based on the center for the entire dataset, which is effectively the reference genome.

**Table 3:** Entropy of all clusterings of GISAID 1 dataset computed^*a*^

| *k*-modes setting<br>(initialization + distance) | without gap filling |  | with gap filling |  |
| --- | --- | --- | --- | --- |
|  | expected<br>entropy | total<br>entropy | expected<br>entropy | total<br>entropy |
| **without clustering** | 9536.89 | 9536.89 | 8417.89 | 8417.89 |
| **random centers + Hamming** | 123.00 | 3170.60 | 109.21 | 2474.30 |
| **random centers + TN-93** | 127.32 | 4401.18 | 111.05 | 3470.03 |
| **pairwise distant + Hamming** | 422.65 | 4651.23 | 294.98 | 3629.47 |
| **pairwise distant + TN-93** | 273.34 | 3500.14 | 256.44 | 3007.07 |
| **CliqueSNV + Hamming** | 110.58 | 2585.29 | 90.42 | 2308.95 |
| **CliqueSNV + TN-93** | 121.87 | 2379.46 | 100.85 | 2117.40 |

<sup>a</sup> The expected entropy (Eq. 4) and total entropy (Eq. 5) of the sequences of the GISAID 1 dataset without clustering (*i.e.*, considered as a single cluster containing all sequences), and when clustering using each of the six combinations of settings mentioned in Sec. 2.2, both without filling gaps and with gap filling.

Table 4 reports the runtimes of the various stages of our CliqueSNV-based clustering approach, and Table 5 compares the overall runtimes of CliqueSNV-based clustering and the *k*-modes approach. We note, given the latter table, that the CliqueSNV-based approach had a slightly lower runtime than the *k*-modes approach, despite it performing best overall.

**Table 4:** Runtime of each stage of the CliqueSNV-based approach^*a*^

| Stage | Time (seconds) |
| --- | --- |
| CliqueSNV (finding initial centers) | 2405.08 |
| clustering (with $k$-modes) | 2324.34 |
| gap filling | 2740.32 |
| entropy computation | 1254.22 |
| **Total** | **8723.96** |

<sup>a</sup> Runtime of each of the different stages of the CliqueSNV-based approach on the GISAID 1 dataset. All stages were executed on a PC with an Intel(R) Xeon(R) CPU X5550 2.67GHz x2 with 8 cores per CPU, DIMM DDR3 1333 MHz RAM 4Gb x12, and running the CentOS 6.4 operating system.

**Table 5:** Runtimes of CliqueSNV-based and *k*-modes clustering^*a*^

| Clustering method | Time (seconds) |
| --- | --- |
| CliqueSNV-based | 4729.42 |
| $k$-modes | 4922.44 |

<sup>a</sup> Runtimes of CliqueSNV-based and $k$ -modes (random centers + Hamming) for the GISAID 1 dataset. Both methods were executed on a PC with an Intel(R) Xeon(R) CPU X5550 2.67GHz x2 with 8 cores per CPU, DIMM DDR3 1333 MHz RAM 4Gb x12, and running the CentOS 6.4 operating system.

### 5.2 Entropy minimization

The main goal of entropy minimization is to make further gains on the performance of existing clustering techniques. Hence, we apply our Monte-Carlo based procedure described in Sec. 2.4 to the clustering obtained by our CliqueSNV-based method (the most performant method), which identified 28 subtypes in the GISAID 1A dataset. As a baseline for comparison, we also produce a random clustering of the data into 28 clusters. Table 6 reports results of our Monte-Carlo based entropy minimization procedure on these two initial clusterings when preprocessing to various different numbers *n* of tags. Initial clustering with our CliqueSNV-based method followed by our Monte-Carlo procedure with 1000 tags achieved the largest decrease in entropy, from 2093.6 to 945.9, as well as the best overall final clustered entropy. The results for CliqueSNV-based clustering indicate that a local (probably global) minimum sits somewhere between 1500 and 500, in terms of the optimal number *n* of tags to select to achieve the best results. While the results for random clustering were considerably worse, there seems to be a trend towards better entropy with reduced numbers of tags. Finally, Fig. 3a and Fig. 3b depict the entropy descent of our Monte-Carlo method applied to the initial CliqueSNV-based clustering for 28000 and 1000 tags, respectively. The latter shows more (relative) improvement in the entropy, indicating that selecting a subset of tags can allow the Monte-Carlo iteration to approach closer to the optimum entropy with fewer iterations.

**Table 6:**
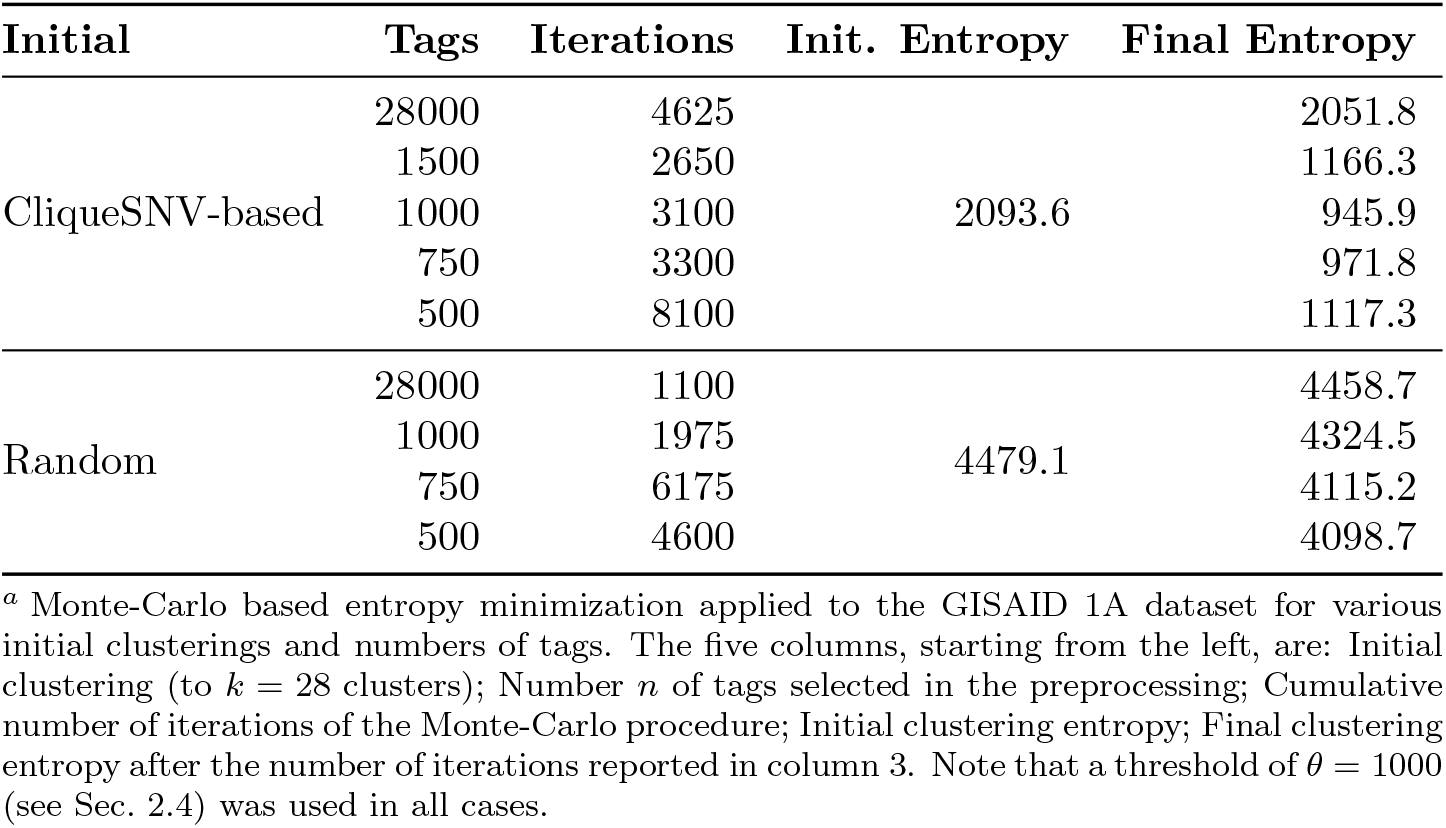
Entropy minimization with the GISAID 1A dataset^*a*^

### 5.3 Finding subtypes

One of the important goals of clustering in this context is to identify subtypes, *e.g*., variants of concern (VoCs), etc. Here we demonstrate the ability of our clustering approaches to finding subtypes in the UK dataset, and then in the much larger GISAID 2 dataset.

#### 5.3.1 The UK dataset

Using the UK dataset, our CliqueSNV-based approach identified 15 subtypes. Since the data here are over a shorter time span (are smaller) and more uniform, a *k* of 15 was sufficient for the minimum cluster frequency to be at least 1% of the population. On the other hand, MeShClust James *et al*. (2018) was only able to find 3 clusters in this data set. Table 7 reports the *F*_1_ score of the methods we compared. Our CliqueSNV-based approach outperformed other methods by a large margin in producing a clustering with all sequences of the Alpha variant residing in a single cluster, while 1.30% of the sequences in this cluster do not belong to the Alpha variant. In the clustering produced by the *k*-modes approach, the sequences of the Alpha variant were spread over five clusters, while one cluster contained 97.45% of the Alpha variant sequences. However, 86.54% of the sequences of this cluster did not belong to the Alpha lineage. MeShClust, on the other hand, produced a clustering with all Alpha variant sequences residing in a single cluster, where 90.68% of the sequences in this cluster do not belong to the Alpha variant.

**Table 7:** *F*_1_ score of various clustering approaches on the UK dataset^*a*^

| Method | $F_1$ score | $F_1$ largest cluster |
| --- | --- | --- |
| CliqueSNV-based | 0.99 | 0.99 |
| $k$-modes | 0.003 | 0.24 |
| MeShClust | 0.11 | 0.11 |

<sup>a</sup> The $F_1$ score of the clustering produced by a method (column 1) with respect to the sequences of the Alpha variant (column 2), and, of the cluster containing the largest number of sequences with the Alpha variant (column 3).

We report the relative distributions of these different subtypes in Fig. 4 over the period of time between the beginning of October 2020, when the first case of the Alpha variant was reported in the UK, to the middle of December, when this variant comprised more than one third of all sequences. We report a weekly moving average because a weekly oscillation in SARS-CoV-2 data has been noted in Bukhari *et al*. (2020). One will notice, in Fig. 4, the sharp increase of the relative proportion of a certain subtype (in red) to more than a third of the population. We confirm from metadata that this corresponds to the Alpha variant that was first identified in studies such as Volz *et al*. (2021b).

**Figure 4:**
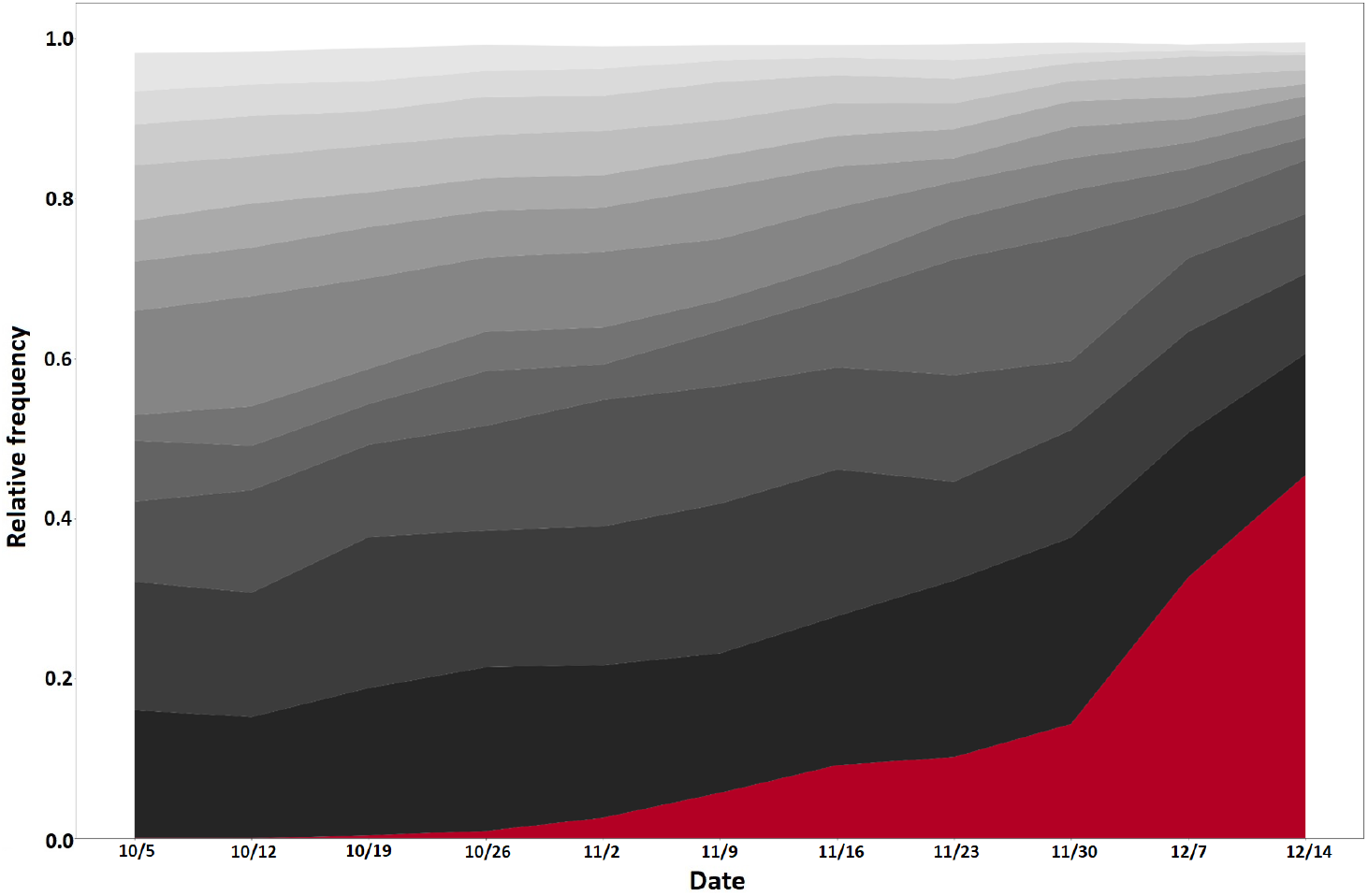
Subtype distribution (the UK dataset, weekly window, relative count), produced our CliqueSNV-based clustering method. The subtype in red contributes to sequences that correspond to the Alpha variant.

When restricting the clusters returned by our CliqueSNV-based approach to the final one-week interval of Fig. 4, leading up to mid December 2020, all Alpha variant sequences appear in a single cluster (among a total of 15). In the clusters returned by the *k*-modes approach, on the other hand, sequences of the Alpha variant are spread over 13 clusters, with counts ranging from 1 to 6327 Alpha variant sequences per cluster. MeShClust again produced only 3 clusters, with a single cluster containing all Alpha variant sequences, when restricted to this final interval, while 90.86% of the sequences in this cluster did not belong to the Alpha variant. The expected entropy of our CliqueSNV-based approach and the *k*-modes approach were 75.73 and 94.16, respectively, while the total entropy was 986.48 and 2074.12, respectively. This illustrates the ability of our clustering to identify subtypes which are known in the literature. Interestingly enough, the study of Volz *et al*. (2021b) is based on an approach of building a phylogenetic tree. This demonstrates our approach, which is based on clustering sequences, as a viable alternative.

Our CliqueSNV-based clustering method was able to detect one subtype which tends to dominate the population in this UK dataset, in attaining good entropy and *F*_1_ scores. However, we wanted to further validate if this is consistent with other independent measures of quality, such as the cluster-based fitness coefficient that we detail in Sec. 3.2. To compute this, we chose our time points *t* to be intervals of one week over the period from the beginning of October to the middle of December, exactly as in Fig. 4. The size *X*_*i*_(*t*) of each cluster *C*_*i*_ (of *k* = 15 clusters) for every week *t* was obtained, and each fitness coefficient *r*_*i*_ was computed according to Eq. 8. In order to reduce sampling error, we drew 2000 random samples from the Poisson distribution on *X*_*i*_(*t*) according to Sec. 3.2. We repeated this 100 times, and we report in Table 8 the 95% confidence interval of the top five clusters, sorted by interval lower bound. We note that similar results are obtained with either Hamming or TN-93 distance, with TN-93 distance corresponding to slightly higher fitness coefficients. We confirm that in either case, the mostly highly ranked cluster in terms of fitness (with cluster ID 6) corresponds to the cluster containing all of the sequences pertaining to the Alpha variant from above. This highlights the ability of our clustering-based approach for detecting, based purely on sequence content, novel subtypes which have the potential of becoming dominant in the population.

**Table 8:**
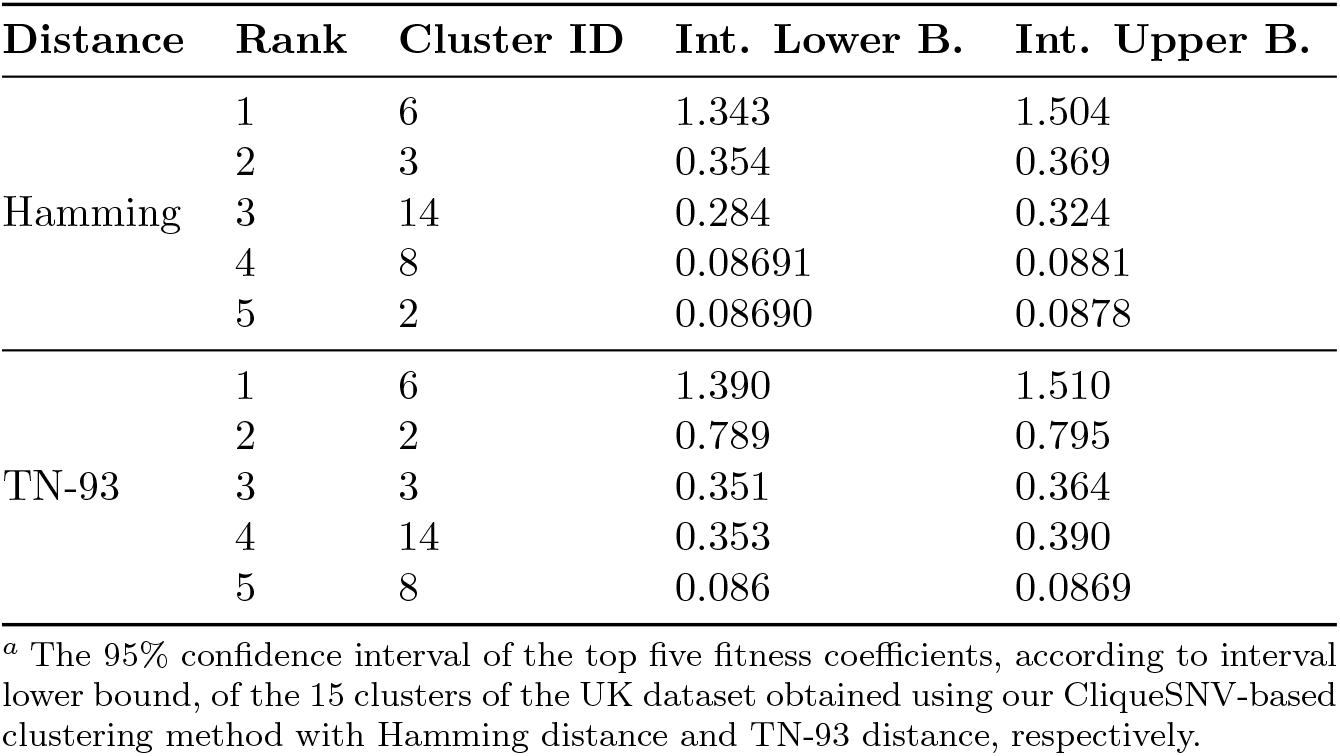
Fitness coefficients of the clusters of the UK dataset^*a*^

| Distance | Rank | Cluster ID | Int. Lower B. | Int. Upper B. |
| --- | --- | --- | --- | --- |
| Hamming | 1 | 6 | 1.343 | 1.504 |
|  | 2 | 3 | 0.354 | 0.369 |
|  | 3 | 14 | 0.284 | 0.324 |
|  | 4 | 8 | 0.08691 | 0.0881 |
|  | 5 | 2 | 0.08690 | 0.0878 |
| TN-93 | 1 | 6 | 1.390 | 1.510 |
|  | 2 | 2 | 0.789 | 0.795 |
|  | 3 | 3 | 0.351 | 0.364 |
|  | 4 | 14 | 0.353 | 0.390 |
|  | 5 | 8 | 0.086 | 0.0869 |

<sup>a</sup> The 95% confidence interval of the top five fitness coefficients, according to interval lower bound, of the 15 clusters of the UK dataset obtained using our CliqueSNV-based clustering method with Hamming distance and TN-93 distance, respectively.

#### 5.3.2 The GISAID 2 dataset

Since our CliqueSNV-based clustering approach was able to clearly pinpoint the Alpha variant within the UK dataset, we tested it also on the GISAID 2 dataset, which contains many of the variants listed in Table 1. CliqueSNV-based clustering identified 36 subtypes in this dataset. We first computed fitness coefficients *r*_*i*_ (Eq 8) for these 36 clusters using one week time intervals *t*. Table 9 reports the 95% confidence interval due to subsampling (see Sec 3.2) of the top and bottom five clusters, sorted by interval lower bound. One will notice immediately that fitness coefficient is much more evenly distributed across the clusters of this dataset, compared to the UK dataset (Table 8).

**Table 9:** Fitness coefficients of the clusters of the GISAID 2 dataset^*a*^

| Rank | Cluster ID | Int. Lower B. | Int. Upper B. |
| --- | --- | --- | --- |
| 1 | 1 | 0.0601 | 0.0602 |
| 2 | 17 | 0.0486 | 0.0489 |
| 3 | 21 | 0.0463 | 0.0463 |
| 4 | 20 | 0.0456 | 0.0457 |
| 5 | 35 | 0.0440 | 0.0440 |
| 32 | 4 | 0.0143 | 0.0143 |
| 33 | 29 | 0.0138 | 0.0138 |
| 34 | 28 | 0.0120 | 0.0120 |
| 35 | 32 | 0.0118 | 0.0118 |
| 36 | 34 | 0.0110 | 0.0110 |

<sup>a</sup> The 95% confidence interval of the top and bottom five fitness coefficients, according to interval lower bound, of the 36 clusters of the GISAID 2 dataset obtained using our CliqueSNV-based clustering method. The mean ( $\mu$ ) ± standard deviation ( $\sigma$ ) of the interval lower and upper bounds are $0.0281 \pm 0.0122$ and $0.0281 \pm 0.0122$ , respectively.

Table 10 reports some the variants found by our CliqueSNV-based approach in terms of specificity, *F*_1_ score and fitness rank (Table 9). Notice that specificity/*F*_1_ score generally decreases with rank and cluster size, as would be expected. Exceptions to this trend are the Gamma/Zeta variant in *F*_1_ score vs. Rank (having a high *F*_1_ score for its rank) and the Epsilon variant (having a large cluster size for its *F*_1_ score and rank). Finally, since the GISAID 2 dataset contains more than 1 million sequences, the Gamma/Zeta, Beta and Epsilon variants comprise less than 1% of the sequences, yet our CliqueSNV-based was still able to identify them with specificities around 50% and *F*_1_ scores *≥* 0.5. This demonstrates the ability of our clustering approach to detect rare subtypes in very large sets of sequences.

**Table 10:** Variants found in the GISAID 2 dataset using CliqueSNV-based clustering^*a*^

| Variant | ID | Specificity | $F_1$ | Rank | Size |
| --- | --- | --- | --- | --- | --- |
| Alpha (UK) | 1 | 93.16% | 0.96 | 1 | 265 255 |
| Gamma & Zeta (Brazil) | 25 | 51.21% | 0.68 | 7 | 1892 |
| Beta (S. Africa) | 21 | 45.85% | 0.62 | 3 | 2754 |
| Epsilon (California) | 13 | 41.08% | 0.58 | 13 | 9251 |

<sup>a</sup> Specificity, $F_1$ score and fitness rank (Table 9) of the cluster containing the largest number of sequences of the corresponding variant.

## 6 Conclusions

In this work, we successfully adapted a method CliqueSNV Knyazev *et al*. (2020), originally designed for discovering viral haplotypes in an intra-host population, to finding subtypes of SARS-CoV-2 in the (massively inter-host) global population. We use clustering entropy Li *et al*. (2004) to assess the quality of a clustering — a notion which naturally reflects the underlying processes from which a set of viral subtypes arises. We introduce two additional techniques which boost the entropy even further, namely, gap filling and Monte Carlo entropy minimization. The former is useful for sequences collected before March 2020 when collection and sequencing were not yet refined, while the latter is possible because clustering entropy is convex Li *et al*. (2004), allowing optimization techniques aimed directly at minimizing entropy as the objective. We show that our CliqueSNV-based clustering method outperforms other techniques in terms of low entropy, and the further improvements in entropy which can be obtained with gap filling and Monte Carlo minimization.

We then turned to datasets obtained from the GISAID and EMBL-EBI (UK) databases in order to identify viral subtypes. Our method was able to most clearly identify the Alpha variant in the UK dataset, with a single cluster containing all sequences with a specificity > 99%. These results tended to be in agreement with the entropies obtained, as well as with the measure of selective fitness introduced in Sec. 3.2. In the GISAID dataset, which contains over one million sequences, our CliqueSNV-based method was able to clearly identify the Alpha variant, but also the lesser represented Beta (South Africa), Epsilon (California), Gamma and Zeta (Brazil) variants. What is interesting about this is that these lesser represented variants comprise a few thousand sequences each (< 1% of the sequences), and yet our method was able to cluster them with specificities around 50%, corroborating again with the fitness coefficient. This demonstrates the approach of clustering as a viable and scalable alternative for detecting even the rarest subtypes at an early stage of development.

An immediate future work is a more full exploration of how our Monte Carlo entropy minimization approach can be made faster and more scalable to large datasets. Ideas include parallelization of our current approach, the design of data structures that can be more efficiently updated, or heuristics beyond our use of tags. The use of optimization techniques other than the Monte Carlo method is a possibility as well. Since CliqueSNV Knyazev *et al*. (2020) is a relatively new technique, possible advancements in its ability to better detect viral haplotypes within an intra-host population would likely carry over to improvements to finding subtypes in the inter-host population setting of this work. Finally, while we provide a viable alternative to building phylogenetic trees (*e.g*., du Plessis *et al*. (2021)) for detecting subtypes, it would be interesting to explore how these could be combined (as in *e.g*., Ciccolella *et al*. (2021a)).

## Acknowledgements

AM, SK, BS, RH and AZ were partially supported from NSF Grant 1564899 and NIH grant 1R01EB025022-01. FM and PS were partially supported from NIH grant 1R01EB025022-01 and NSF Grant 2047828. AM, BS and SK were partially supported by GSU Molecular Basis of Disease Fellowship. MP was supported by a Georgia State University / Computer Science startup grant.

